# Early Jurassic origin of avian endothermy and thermophysiological diversity in Dinosauria

**DOI:** 10.1101/2023.12.21.572807

**Authors:** Alfio Alessandro Chiarenza, Juan L. Cantalapiedra, Lewis A. Jones, Sara Gamboa, Sofía Galván, Alexander J. Farnsworth, Paul J. Valdes, Graciela Sotelo, Sara Varela

## Abstract

A fundamental question in dinosaur evolution is how they adapted to substantial long-term shifts in Earth System during the Mesozoic and when they developed environmentally independent, avian-style acclimatization due to the evolution of an endothermic physiology. Combining fossil occurrences with macroevolutionary and paleoclimatic models, we unveil distinct evolutionary pathways in the main dinosaur lineages: ornithischians and theropods diversified across broader climatic landscapes, trending toward cooler niches. An Early Jurassic shift to colder climates in Theropoda suggests an early adoption of endothermic thermophysiology. Conversely, sauropodomorphs exhibited prolonged climatic conservatism associated with higher thermal conditions. Paleo-biome mapping emphasizes temperature, rather than plant productivity, as the primary driver of this pattern, suggesting poikilothermic physiology with a stronger dependence on higher temperatures in sauropods since the Early Jurassic.

**One-Sentence Summary:** Dinosaur climatic evolution reveals early endothermy emergence in theropods, ornithischians but heterotherm sauropodomorphs.

---

Dinosaurs ubiquitously occupied the Earth’s terrestrial ecosystems for over 160 million years (from 230 to 66 million years ago) (*1*). These animals evolved and diversified into a remarkable array of ecological niches, adopting diverse dietary preferences and lifestyles (*1*, *2*), and unfolding into an impressive body size spectrum spanning eight orders of magnitude (*3*): from 80- ton sauropods (*4*), which stand as the largest land animals, to some contemporary hummingbirds weighting just a few grams. Over the past two centuries, while illuminating the ancestral connection between dinosaurs and modern birds (*5*, *6*), extensive research has unveiled a level of ecomorphological diversity within dinosaurs that rivals many other terrestrial vertebrate groups.

Accumulating evidence on dinosaur paleobiology (*7*–*9*) indicates that the development of endothermy (the ability to self-regulate body temperature) may have been a pivotal factor in their ecological diversification (*10*). Endotherm, warm-blooded dinosaurs could have flourished into harsher environments, including high-latitude cold regions (*11*, *12*). This view raises intriguing questions about the origins of key biological innovations once considered exclusive to neornithine birds but now recognized as emerging within this ancient lineage of stem birds (*6*). The conventional perception of Dinosauria regarded these animals as a group of stereotypically lumbering, slow-moving reptiles, sharing characteristics with most ectothermic, heterothermic, and poikilothermic vertebrates. That is, species with higher variation in internal temperature that rely on environmental heat for homeostasis (*7*). However, recent discoveries [(*12*, *13*)] are challenging this ‘cold-blooded’ environmental, thermally-regulated body temperature paradigm, painting a more nuanced picture. It highlights the emergence – at least within certain dinosaurian subclades (e.g. Maniraptora) – of traits typical of endothermic, homeothermic, and tachymetabolic (highly active) tetrapods capable of maintaining a constant body temperature through internally generated heat (*14*).

While this view is gaining traction, it is essential to acknowledge that not all facets of the scientific community uniformly accept it (*15*–*20*). Various lines of evidence, including comparative anatomy (*21*), reproductive paleobiology (*22*), ecology and energetics (*23*), biomechanics (*23*), osteohistology (*24*), paleobiogeography (*17*), isotopic geochemistry (*25*, *26*), and soft tissues (*19*, *20*, *27*) show at best loose consilience, all implying: 1) conflicting views on the specific thermophysiological strategies of different dinosaur subclades, and 2) consequent lack of consensus in the timing and modality of the appearance of endothermy in the stem-avian lineage. One corollary of the differential thermophysiological strategies among terrestrial tetrapods is the potential for endotherms to expand their latitudinal range, owing to their reduced reliance on environmental temperature (*28*). For instance, modern birds and mammals have achieved nearly global distributions, extending from the tropics to polar regions. In contrast, most reptilian lineages, such as squamates, turtles, crocodilians, and amphibians, primarily inhabit regions closer to the tropics, their distribution and diversity predominantly constrained by temperature, except for well-defined hyperthermal events in the geological past (*29*). Determining when this broader macroecological pattern emerged within the avian lineage remains a pivotal question, strongly contingent upon uncovering the origin of these critical physiological traits.

To overcome the limitations of a rigid typological rationale, often imposing strict ontological boundaries on continuous systematic groups (*30*), we adopt an evolutionary perspective to gain fresh insights into the diverse thermophysiological strategies exhibited by dinosaurs. By integrating concepts of *tempo* and *modo* from established macroevolutionary theory in vertebrate paleontology (*31*), our approach employs a phylogenetic comparative method framework. This apporach quantitatively explores evolutionary patterns signaling likely shifts in thermophysiology due to the invasion into novel adaptive landscapes (*sensu* Simpson (*31*)), and specifically in terms of climatic tolerance. Through the application of phylogenetic comparative methods and global paleoclimate modelling tools (*32*), including climatically calibrated dinosaur phylogenies analysed through a set of evolutionary models (*33*–*35*), we reconstructed the climatic adaptive landscape (*36*) of Dinosauria throughout the Mesozoic.

We tested three hypothetical scenarios for the evolution of the dinosaurian climatic niche landscape: 1) relaxed plasticity, involving a continuous invasion of new climatic niches without directional limitations imposed by physical factors; 2) the influence of internal (phylogenetic or clade-dependent) constraints; and 3) the impact of external (climatic regime shifts) factors at specific time horizons. Evolution shaped by external pressures may have influenced climatic occupation in the adaptive landscape, prompting biotic adaptations in terms of thermophysiological plasticity or the differential success of climatically more adaptive lineages. Additionally, we examined how biome occupation during the Mesozoic influenced the exploitation of suitable climates for dinosaurs, potentially shaping our macroevolutionary interpretations. Our study directly assessed the timing and modes of transitions to different climatic niches in dinosaur evolution, exploring significant climatic events spanning 160 million years that affected dinosaurian climatic niches. This allowed us to infer distinct thermophysiological strategies within different Dinosauria subclades, considering how Mesozoic climatic zone occupation constrained dinosaurian climatic history and shaped our macroevolutionary interpretations.

## Results

### The occupation of climatic niche space throughout the Mesozoic

The evolution of dinosaurian climatic niches followed two main dimensions (Fig. 1), one (principal component 1, PC1) controlled by a combination of maximum and minimum temperatures, minimum precipitation, and precipitation seasonality and the other (PC2) by minimum temperature and precipitation seasonality. Dinosauromorphs started diversifying in dry environments characterised by high temperatures (Fig. 1A). From these arid conditions, sauropodomorphs first expanded throughout this drier and warm niche, stalling at high temperatures, but migrating towards more seasonal precipitation conditions in later diverging groups. Ornithischians followed cooler and wetter environments through their evolution, with theropods tracking the other dinosaur subclades. Although both theropods and ornithischians evolved towards lower temperatures (Fig. 1B), Ornithischia exhibited a more pronounced preference for cooler conditions, especially in later thyreophorans, ceratopsians, and hadrosaurids. Earlier diverging neornithischians occupied warmer and more seasonal conditions, closer to their dinosauromorph relatives. Theropods displayed a wide climatic niche, invading colder but slightly more seasonal precipitation conditions in later diverging carnosaurs and coelurosaurs (including avialians), while still maintaining ancestrally warmer and drier niches in some members of all these clades. Overall, ornithischians and theropods increased their climatic niche disparity compared to sauropodomorphs and ancestral dinosauromorphs. Ornithischia and Theropoda were both projected mainly towards cooler niches, with a preference for more humid conditions in the former and higher seasonality (in terms of humidity) in the latter (Fig. 1B).

**Figure 1.**
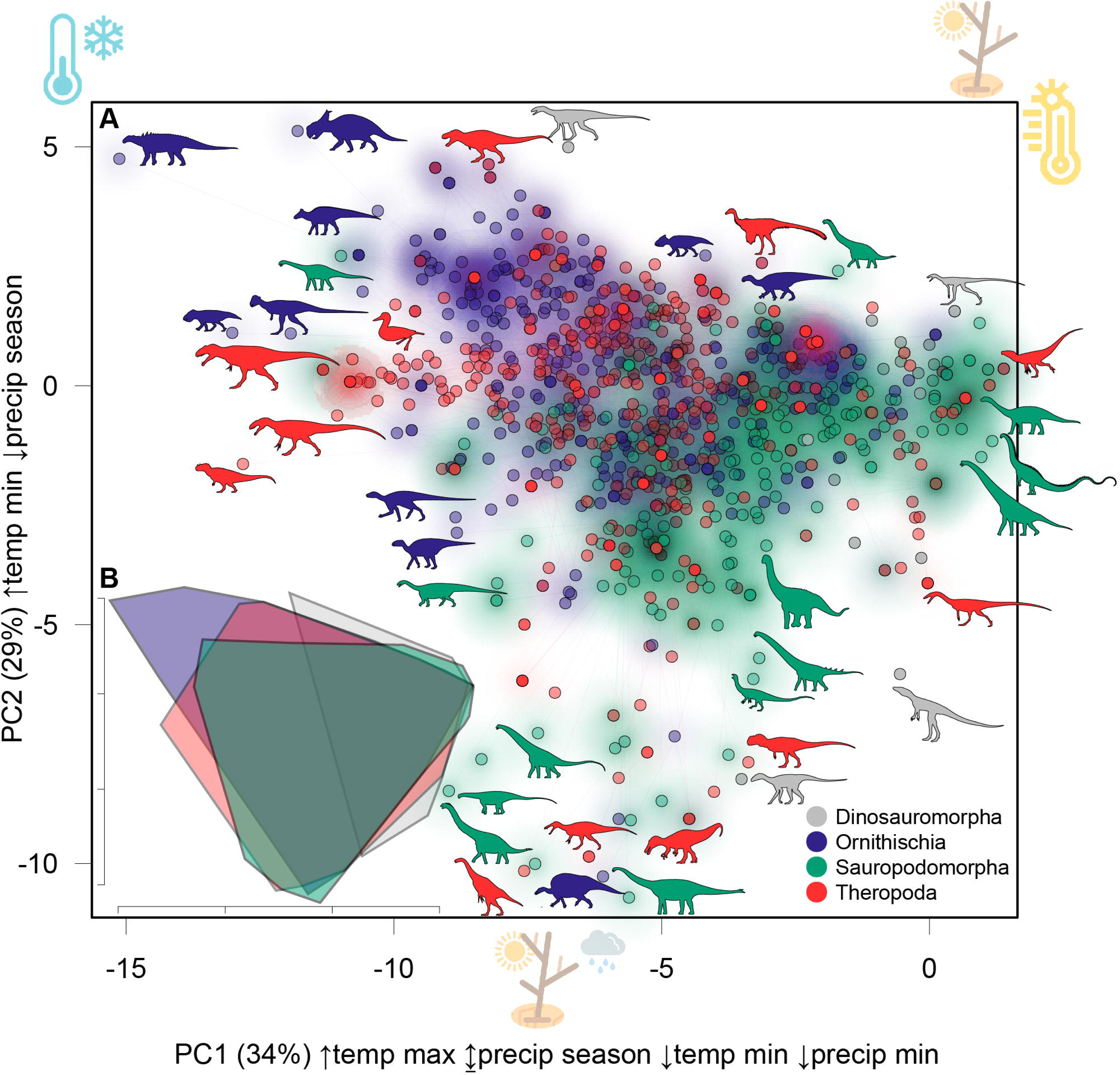
Phylogenetic-climatic niche space for Mesozoic dinosaurs. A phylogenetic Principal Component Analysis (PCA) (**A**) represents the projection of the Dinosauria supertree (see Methods) into a PCA of climatic variables. PC1 axis shows strong positive correlation with maximum temperature (↑temp max), low positive correlation with precipitation seasonality (↕precip season), strong negative correlation with minimum temperature (↓temp min), and strong negative correlation with minimum precipitation (↓precip min). PC2 axis shows strong positive correlation with minimum temperature (↑temp min) and negative correlation with precipitation seasonality (↓precip season). Shadows around points highlight the relative density in the principal component space of respectively non dinosaurian Dinosauromorpha (gray), Ornithischia (blue), Sauropodomorpha (green) and Theropoda (red). Lower left plot (**B**) shows 95% confidence interval convex hulls for each dinosauromorph subclade. Blue thermometer (top left corner) symbolizes the direction of the vector in the PC space region for cold temperatures; yellow thermometer (top right corner) indicates the direction of the vector in PC space for warm temperatures; brown shrub (top right corner) depicts dry conditions, while the same with a gray, rainy cloud (mid, lower side of the graph) illustrate seasonal conditions. Gray silhouettes (**A)** depict Dinosauromorpha, blue Ornithischia, green Sauropodomorpha and red Theropoda.

### Macroevolution of the climatic adaptive landscape in Dinosauria

The evolution of climatic niches in Mesozoic dinosaurs is expressed differentially among the main subclades of Dinosauria (Fig. 2). Sauropodomorphs are restricted to higher temperatures (25.4–23.3°C), while ornithischians and theropods exhibit broader thermal ranges. Overall, we find that no single regime model can explain the entire thermal evolution for each subclade. Instead, the Ornstein-Uhlenbeck models of evolution (OUMA and OUMVA) are favored (Table S1, Data S3), suggesting that climatic niche exploration is influenced by physically constrained models following distinct macroevolutionary thermal optima (θ). For sauropodomorphs (Fig. 2), the model-fit results indicate a conservative evolution from an ancestral temperature of 25.4°C, with an evolutionary rate (σ^2^) at the root of 1.85×10^−5^ and an attraction value (α) of 0.6×10^−1^. Different subclades within Sauropodomorpha reached distinct but close thermal optima, with Neosauropoda reaching an optimum of θ=24.7°C (Fig. 2A) with an almost equivalent attraction value (α) to the root of 0.7×10^−1^ but a slower rate (σ^2^) of 3.2×10^−6^. A rate increase (σ^2^=1.2×10^−5^) at an attraction value (α) of 0.6×10^−1^ leads to a temperature optimum of θ=23.3°C (Fig. 2A) in Somphospondyli (later diverging titanosaurs). A significant transition (Table S1) occurs at the ‘Jenkyns’ early Toarcian Oceanic Anoxic Event (OAE) with a rate increase from σ^2^=0.26×10^−6^ to σ^2^=0.13×10^−3^ with almost equivalent attraction optima (α_1–2_=0.91×10^−1^–0.69×10^−1^) for a temperature decrease from the root state (z_0_) of 28.1°C to a later optimum of θ=23.4°C (Fig. 2A).

**Figure 2.**
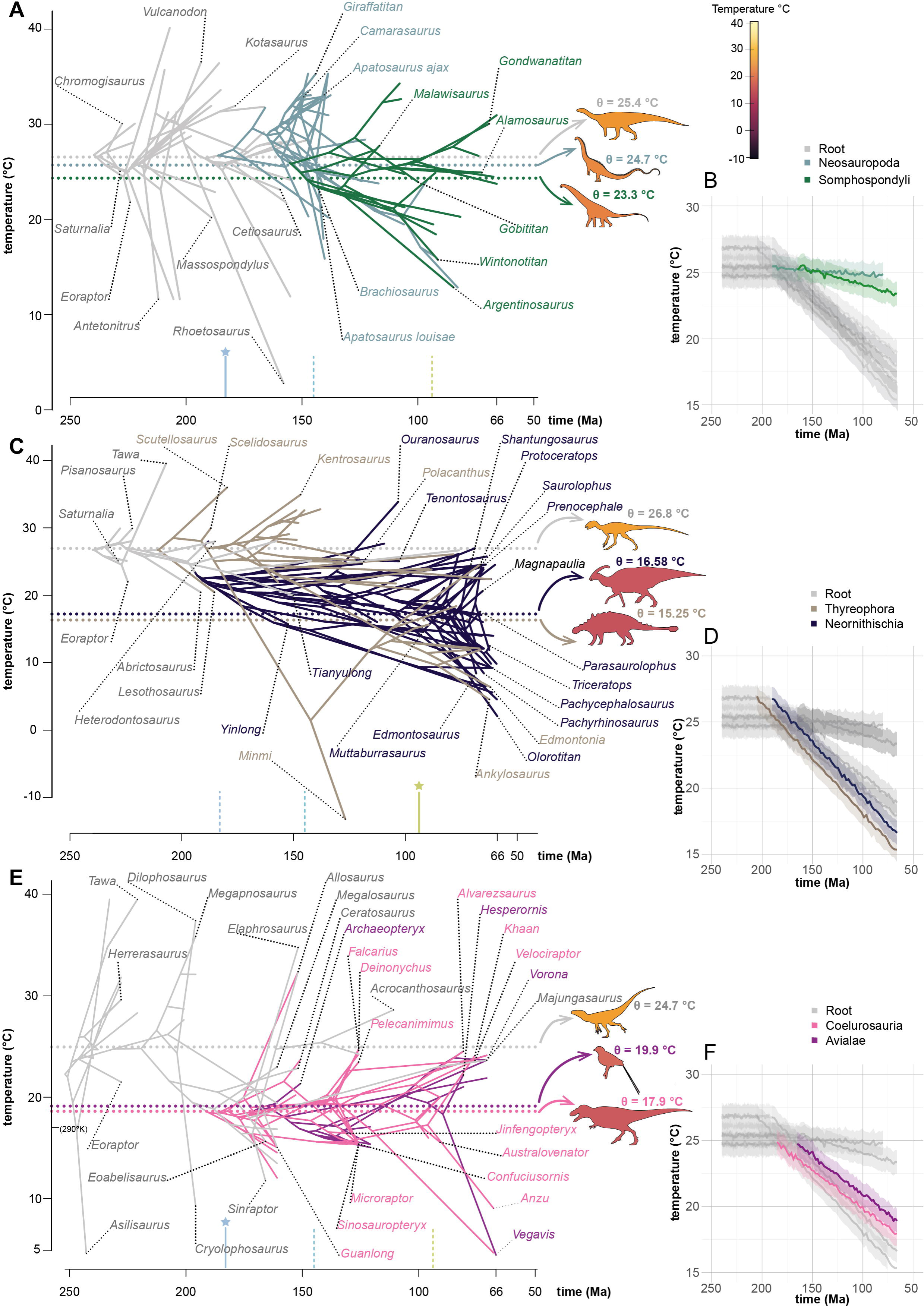
Macroevolutionary climatic landscape in Dinosauria throughout the Mesozoic. Evolutionary regimes along the temperature axis (in °C) are shown for Sauropodomorpha (**A**, **B**), Ornithischia (**C**, **D**) and Theropoda (**E**, **F**). Clade partitions along the trees (**A**, **C**, **E**) are portrayed in different colors with horizontal dotted lines showing optima (θ) for each clade partitions. Trends in temperature optima occupation are simplified in **B**, **D**, **F**. Dinosaur silhouettes colors change according to the temperature at each θ. Stars on top of vertical bars indicate significant (following model choice according to AIC values; see Methods; Table S1) changes in time-dependent partitions across the ‘Jenkyns’ event (∼183 Ma), the Jurassic/Cretaceous (∼145 Ma) and the Cenomanian/Turonian (93.9 Ma) boundaries (with dotted lines showing non-significant trend variations). Silhouettes were obtained from http://phylopic.org/ (see Acknowledgements).

Ornithischia undergoes a rapid transition (σ^2^=0.59×10^−7^ to σ^2^=0.14×10^−4^) from a warm climate niche starting at z_0_=26.8°C in the Early Jurassic, to a cooler climatic niche in Thyreophora, reaching θ=15.25°C, accompanied by a lower attraction value (α=1.5×10^−2^; Fig. 2C, D). In Neornithischia (Fig. 2C), the rate slows down (σ^2^=0.54×10^−5^), while the attraction to a comparatively cooler niche of θ=16.58°C increases (α=3.7×10^−2^). The regime transition is sub-optimally modeled to occur (Table S1) at the Cenomanian/Turonian (C/T) boundary (Fig. 2C), with negligibly different attraction values (α_1–2_=0.4×10^−1^ – 0.3×10^−1^) from pre-C/T values. The favored clade-partitioned model has equal attraction and rate values (OUMA), depicting a transition from z_0_=26.8°C to θ=15.25°C in Thyreophora and θ=16.58°C in Neornithischia (Fig. 2C).

Theropods (Fig. 2E, F) exhibit an abrupt decrease in temperature optima favored in the model partitioned between their root, Coelurosauria, and Avialae, compared to the stepwise pattern when partitions are modeled at the base of the dinosauromorph node, at the base of Theropoda, and in Coelurosauria. This model displays a decrease in rates (σ^2^=0.1×10^−4^ to σ^2^=0.3×10^−6^) with a slight increase in attraction values (α_1–2_=0.37×10^−1^–0.64×10^−1^), from their root temperature optimum of z_0_=24.7°C to a lower optimum of θ=19.9°C in Coelurosauria. Avialae (Fig. 2E) experiences an acceleration in rates (σ^2^=0.5×10^−4^) with a slight decrease in attraction values (α=0.25), resulting in a temperature optimum of θ=17.9°C. The ‘Jenkyns’ (Fig. 2E) is the only temporal event playing a role in partition transitions (Table S1), with changes in rates (σ^2^=0.45×10^−4^ to σ^2^=0.2×10^−6^), attraction values (α_1–2_=0.13×10^−1^–0.19×10^−1^), and cooling in temperature optima from z_0_=28.03 °C to θ=18.82 °C.

### Testing dinosaurian occupation of climatic zones due to biome preferences

Sauropodomorph evolution through warm niche landscapes (Thermal Bound Hypothesis; TBH) could be alternatively interpreted as a secondary by-effect of their large size: large terrestrial primary consumers preferentially inhabit regions of elevated productivity (*37*), hence they might have preferred lower latitudes, the Productivity Bound Hypothesis (PBH). A quantification on whether Jurassic–Cretaceous sauropodomorphs are bounded to low latitude due to their selective or exclusive occupancy of tropical, highly productive biome is currently lacking. To test these hypotheses (i.e. linking sauropodomorph distribution to biomes) we plotted the taxa used in our phylogenetic modelling analyses on paleogeographic maps (Fig. S12-S14) of broad climatic zones according to an adapted Köppen scheme (*38*, *39*). Recognizing the substantial influence of largely unknown factors (such as substrate lithology and taxonomic composition of vegetation) during the Mesozoic, we opted for climatic zones (*40*) over strict biomes, with the assumption that climatic zones could effectively serve as proxies for the latter. The distribution of sauropodomorphs (Fig. 3A) spans from tropical to cold regions, with an equally higher prevalence in tropical, temperate, and arid regions, compared to cold ones and are absent in polar regions. Contrary to the expectation of the PBH, sauropodomorphs are comparatively less represented in tropical regions than ornithischians and theropods, having a relatively higher prevalence than ornithischians in low productivity arid regions, particularly during the Late Jurassic–Early Cretaceous interval. In contrast, Ornithischia, which are highly abundant in tropical zones, are equally diverse in cold and polar zones.

**Figure 3.**
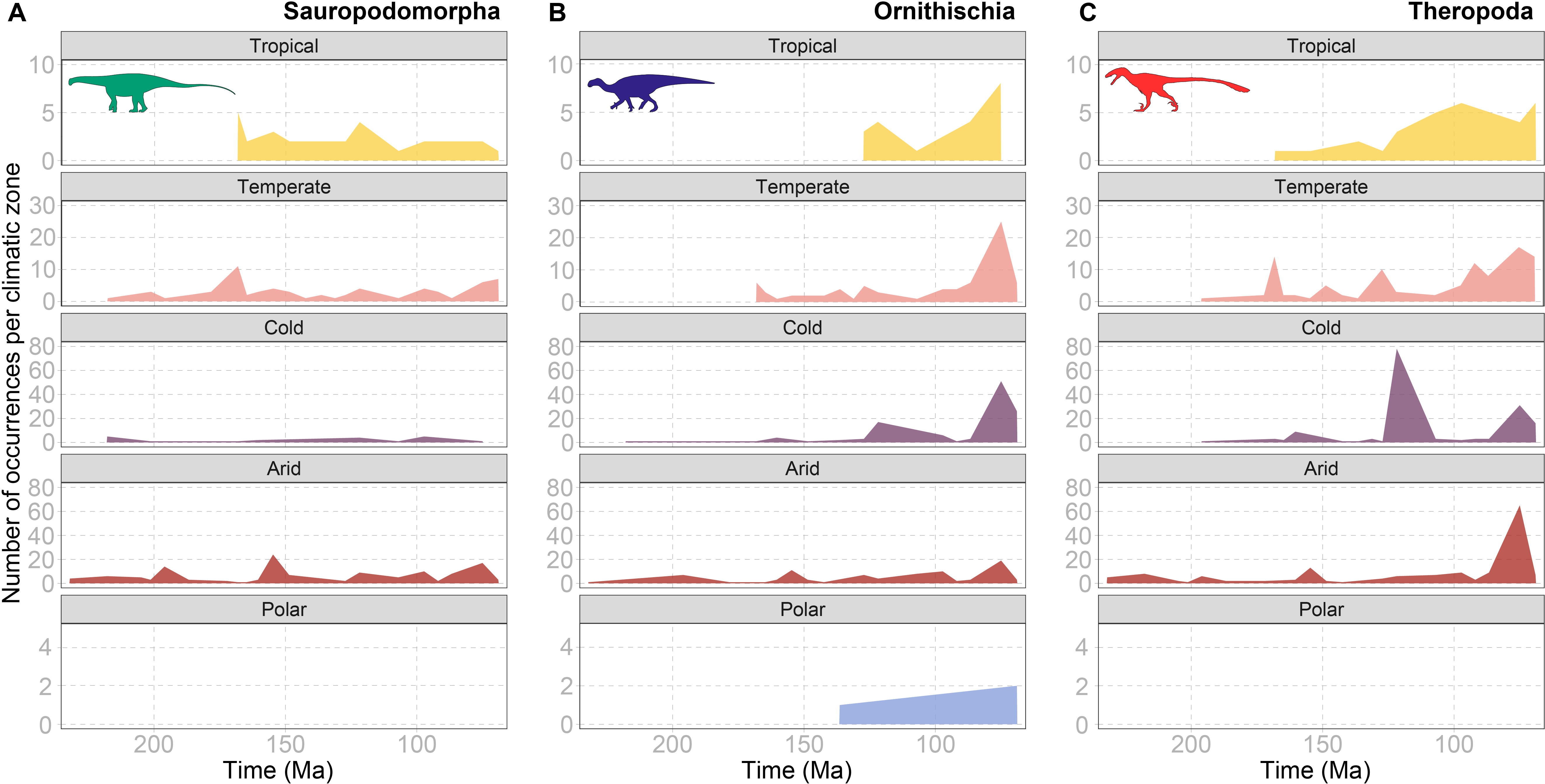
Dinosaur abundance in climatic zones. Taxic-occurrence quantification for Sauropodomorpha (A), Ornithischia (B) and Theropoda (C) in which highest productive climatic zones are arranged on top and less productive ones at the bottom of the figure. Silhouettes were obtained from http://phylopic.org/ (see Acknowledgements).

These observations are consilient with the fossil record of sauropod-rich areas such as the semi-arid and seasonal Late Jurassic Morrison Formation (*17*, *41*) and the arid to seasonal paleoenvironment of the Late Cretaceous Huincul Formation in Argentina (*42*), both sauropod-bearing units yielding some of the largest representative of this clade. These formations have been generated in warm, xeric to seasonal paleoenvironments, conditions preferentially suitable for sauropods (*17*), an observation withstanding potential sampling biases (*17*, *43*, *44*). Furthermore, high latitudes remained highly productive areas in the Mesozoic (*45*, *46*), particularly in the Cretaceous, when pinoid conifers dominated mid-to-high latitudes (*47*). Forested covers thrived in the poles during the Mesozoic (based on evidence from Antarctica and Alaska, especially in Late Cretaceous deposits (*47*–*50*)), hardly exerting a productivity-dependent constrain on dinosaurian primary consumers, as also evidenced by the high-latitude occupation and sustained polar diversity of ornithischian dinosaurs (Fig. 3B).

## Discussion

### Dinosaur diversification throughout the climatic adaptive landscape

The evolution of dinosaur climatic preferences, as represented in multivariate space (Fig. 1), underscores a heightened reliance on temperature-related factors. Early dinosauromorphs appeared and diversified in dry and hot environments, while later diverging dinosaur subclades evolved towards broader niche dimensions, reflecting diverse climatic preferences and thermophysiological constraints. Sauropodomorphs exhibit a preference for high temperatures, while ornithischians and theropods demonstrate a broader range of thermal landscapes in their evolutionary histories. This divergence in climatic niches likely played a crucial role in shaping the ecological diversity, biogeographic history and success of these groups (*12*, *17*, *51*, *52*).

The preference for cooler and wetter conditions within ornithischians, particularly in later thyreophorans, ceratopsians, and hadrosaurids, suggests a trend toward increased humidity and seasonality in their habitats. This adaptation, along with elevated cold tolerance, may have facilitated their continuous persistence at high latitudes and occupation of broader climatic zones (Fig. 3B), ranging from tropical to polar environments (*12*, *17*, *53*). Theropods also displayed a wide range of climatic preferences, with some taxa retaining ancestrally warmer and drier niches, while others adapted to cooler and slightly more seasonal conditions. The temporal-shift phylogenetic models suggest a possible transition towards broader thermal diffusion since the late Early Jurassic ‘Jenkyns’ event, a time likely associated with the radiation of the main tetanuran clades (*54*), and possibly early pannaraptorans (*3*, *55*, *56*).

Our analysis highlights key changes in climatic preferences during specific geological events, such as the ‘Jenkyns’ in the Early Jurassic and the Cretaceous Thermal Maximum (*57*), indicating critical periods of adaptation coinciding with hyperthermals (*58*). The ‘Jenkyns’ event, associated with the Early Toarcian Oceanic Anoxic Event, significantly influenced temperature optima shifts in sauropodomorphs and theropods. Recent empirical evidence (*59*) suggests that eusauropod sauropodomorphs attained large (> 10 m in length) sizes and radiated around this time horizon (180–184 Ma), coinciding with a short-lived episode of global warming, marine anoxia, and a large-scale magmatic event. Concurrent climate perturbations marked a drastic decrease in floral diversity, documented by the contemporaneous rise of conifers. Such an episode might have strongly affected clades of terrestrial animals attaining gigantic size at the time (*59*, *60*), potentially driving the explosion in the diversification of main dinosaurian clades (*58*).

Our results indicate that the exploration of thermal niches during this pivotal transition may have been crucial to dinosaurian success thereafter, due to the consequent ecosystem reorganization and increase in bioregional suitability for the thermically unconstrained ornithischians and theropods (*17*). When the likely more eurythermal (thermally more adaptable) and widely distributed non-eusauropod sauropodomorphs declined (*10*, *61*), the thermophilic eusauropods colonized more tropical latitudes and southern continents (*17*, *52*). Our interpretations reconstruct an early migration of genasaurian (Neornithischia+Thyreophora) ornithischians toward cooler niches, with a stronger temporal signal later at the Cenomanian/Turonian boundary. The early invasion of cool niches by thyreophorans and neornithischians aligns with the current mosaic of morphological traits (from integument (*62*) to ventilation (*63*, *64*)), suggesting an early attainment of a homeothermic (possibly endothermic) physiology in these clades, enabling them to colonize and persist in even extreme latitudes since the Early Jurassic. The strong impact on partition transitions detected for ornithischians at the Cenomanian/Turonian boundary (93.9 Ma) coincides with a hyperthermal event known as the Cretaceous Thermal Maximum (*57*). This event aligns with documented eustatic and atmospheric changes that may have influenced paleodiversity and faunal biogeography (*65*, *66*). This transition from the Early to Late Cretaceous witnessed the evolutionary appearance and radiation of Hadrosauridae (*67*, *68*), highly represented in our dataset. The extent to which the thermal upheaval directly influenced ornithischians (and dinosaurs as a whole) or if the subsequent floral and broader biotic changes had a stronger impact should be the focus of more detailed studies across the Cenomanian/Turonian.

Ornithischians and theropods, the two dinosaurian clades in which protofeathered integument structures have been found (*27*), demonstrate remarkable adaptability across varied climatic zones, while the more thermophilic sauropodomorphs show a specialized preference for low latitudes (*17*, *51*). Notably, these results provide novel insights into the origin of avian endothermy, suggesting that this evolutionary trajectory within theropods towards thermal niche relaxation likely started in the latest Early Jurassic (Fig. 2C), a crucial period for the radiation of coelurosaurs and possibly early diverging avian clades (*56*, *69*, *70*). These findings highlight a potential link between climatic dynamics and the early development of endothermic traits, offering clues to the origin of birds and the heterogenous thermophysiological strategies adopted by Mesozoic dinosaurs. Our study challenges previous notions of static, environmentally conserved dinosaurs, revealing their dynamic, heterogeneous adaptation to diverse climatic conditions throughout the Mesozoic era, contributing to a more nuanced understanding of dinosaurian evolution and the interplay between climate and ecological diversity.

## Acknowledgments

Phil Mannion (UCL) for discussions on sauropod paleobiology. Oliver Rauhut (LMU) for discussions on theropod phylogenies. Silhouettes used in Figure 1 represent the following taxa (clockwise from the higher left corner): *Minmi*, *Edmontosaurus*, *Pachyrhinosaurus*, *Tyrannosaurus*, *Asilisaurus*, *Graciliraptor*, *Harpymimus*, *Altirhinus*, *Gobititan*, *Suzhousaurus*, *Marasuchus*, *Pampadromaeus*, *Herrerasaurus*, *Vulcanodon*, *Diplodocus*, *Giraffatitan*, *Coelophysis*, *Dromomeron*, *Gondwanatitan*, *Tapuiasaurus*, *Anchisaurus*, *Siamotyrannus*, *Diodorus*, *Suchomimus*, *Phuwiangosaurus*, *Ouranosaurus*, *Irritator*, *Tangvayosaurus*, *Nanshiungosaurus*, *Aeolosaurus*, *Rebbachisaurus*, *Chuxiongosaurus*, *Tethyshadros*, *Koreanosaurus*. *Genyodectes*, *Mapusaurus*, *Vegavis*, *Goyocephale*, *Rhoetosaurus*. Silhouettes in Figure 2 (top to bottom) represent the taxa: *Plateosaurus*, *Diplodocus*, *Alamosaurus*, *Heterodontosaurus, Euoplocephalus, Parasaurolophus, Herrerasaurus, Protopteryx* and *Gorgosaurus.* Silhouettes in Figure 3 (left to right) represent the taxa: *Nigersaurus*, *Iguanodon* and *Deinonychus.* All silhouettes throughout this paper are sourced from http://phylopic.org/ and are licensed under Creative Commons licenses: CC BY 3.0 (https://creativecommons.org/licenses/by/3.0/), CC BY-SA.3.0 (https://creativecommons.org/licenses/by-sa/3.0/), and CC BY-NC-SA.3.0 (https://creativecommons.org/licenses/by-nc-sa/3.0/).

## Author contributions

Conceptualization: AAC

Methodology: AAC, JLC, AF, PJV

GCM simulations: PJV, AF

Investigation: AAC, JLC, LAJ, AF, SaG, SoG

Visualization: AAC, SaG, SoG

Supervision: AAC, SV

Writing – original draft: AAC

Writing – review & editing: all authors

## Data and materials availability

quantitative and statistical routines are reported in the Materials and Methods section; fossil datasets (Data S1 to S3) are available on FigShare (a link will be provided upon acceptance, and it is currently included only in the cover letter for the Editor and the Reviewers). Plate models are freely available at https://www.earthbyte.org/. Climate model data can be accessed from the public repository at www.bridge.bris.ac.uk/resources/simulations. All other data are available in the main text, methods, or the supplementary materials.

## Materials and Methods

### Fossil dataset

We developed a specimen-based occurrence dataset, building on previous efforts from Benson et al. (*4*). The occurrences were originally sourced from the Paleobiology Database (paleobiodb.org) but required vetting to ensure that the relevant specimens were correctly anchored to the respective taxa used in the tree and along with the correct stratigraphic and geographic information (Data S1). We modified the dataset for a total of 993 occurrences, updating entries according to current taxonomic information and providing references for previously unavailable or ambiguous data (e.g. locality data). Paleogeographic reconstructions were performed using the Global Plate Model PALEOMAP (*71*), implemented via the ‘paleorotate’ function in the R package paleoverse ver. 1.2.1 (*72*).

### Paleoclimate model

The global paleoclimatic and land-surface model outputs used in this study spans from the Late Triassic (Carnian) to the end-Cretaceous (Maastrichtian). These data were produced from an updated version of the fully coupled Atmosphere-Ocean General Circulation Model (AOGCM) HadCM3, specifically, HadCM3L-M2.1D, following the nomenclature of Valdes, et al. (*73*), developed by the BRIDGE Group (http://www.bridge.bris.ac.uk/resources/simulations). The model has a further critical update that includes modification to cloud condensation nuclei density and cloud droplet effective radius following recent work (*74*–*76*). This raises higher latitude temperatures without significantly changing tropical temperatures reducing the pole-to-equator temperature gradient in line with proxy observations, a persistent problem that has afflicted many paleoclimate models for decades. This update is also found to work under hot, cool, and icehouse climatic conditions, as well as under pre-industrial boundary conditions making it appropriate for use across modern and deep time evolving conditions.

Variables included output by in the general circulation model (GCM) include near-surface (1.5 m) mean annual temperature (°C), near-surface (1.5 m) annual temperature standard deviation (°C), annual average precipitation (mm), annual precipitation standard deviation (mm), net primary productivity (NPP, g C m-2 yr-1), and five plant functional types (broadleaf trees, deciduous trees, shrubs, C3-type and C4-type grasses) using an interactive vegetation scheme called TRIFFID (Top-Down Representation of Interactive Foliage and Flora Including Dynamics) using the MOSES 2.1 land surface scheme at a spatial resolution of 2.75°×3.25° latitude by longitude 278 km by 417 km grid square at the equator). There are 19 hybrid levels in the atmosphere and 20 vertical levels in the ocean with equations solved on the Arakawa B-grid. As is common in all climate models, sub-grid scale processes such as cloud, convection and oceanic eddies are parameterized as they cannot be resolved at the scales required (usually meters to several kilometres) of the model resolution. The ocean model is based on the model of Cox, et al.(*77*) and is a full primitive equation, three-dimensional model of the ocean.

The model simulations were conducted in a similar vein as comprehensively described in Lunt et al. (*78*), Valdes et al. (*79*), and Farnsworth et al. (*80*). The implications of deep-time studies of these GCM constraints were discussed in previous studies (*17*, *29*, *81*–*83*). In summary, the model simulations ran for at least 6000 (often many thousands of years more) model years until they reached full equilibrium in both the atmosphere and deep ocean. Equilibrium is determined by passing three criteria: i) the globally and volume-integrated annual mean ocean temperature trend is less than 1°C per 1000 years; ii) trends in surface air temperature are less than 0.3°C per 1000 years and; iii) net energy balance at the top of the atmosphere, averaged over a 100-year period at the end of the simulation, is less than 0.25/WDm^2^. The variables used in our study represent an annual average of the final 100 years of these simulations. Notably, these models captured temporal fluctuations, regional nuances, and large-scale circulations, including associated energy and momentum fluxes (*78*). Despite the inherent uncertainty in the data, these models effectively replicated the modern-day climates of most terrestrial biomes (*79*). Importantly, HadCM3L played a pivotal role in the Coupled Model Intercomparison Project experiments and demonstrated its utility in various Mesozoic paleobiogeographical investigations (*17*, *29*, *81*, *82*, *84*). The paleogeographic data employed in this study were derived from the dataset created by Scotese & Wright (*71*). Originally conceived as a paleo-digital elevation model (DEM) with a 1°×1° grid, these data were upscaled to match the resolution of the HadCM3L Earth System model (2.75°×3.25°). This upscaling process ensured that the topographic and bathymetric information was broadly preserved, despite its resolution being lowered (*17*, *81*, *85*). These 117 paleogeographic maps (covering the whole of the Phanerozoic) have served as a global atlas, facilitating regional-scale interpretations of paleogeography over the past 540 million years, thereby elucidating the shifting distributions of Earth’s oceans and continents. Here we focus only on 230 to 66 million years ago. Additional information regarding these datasets can be freely accessed at https://www.earthbyte.org/. Land ice was not included within the Scotese paleogeography and is instead transformed onto the model grid assuming a simple parabolic shape to estimate the ice sheet height (m). ‘Realistic’ *p*CO_2_ concentrations for each simulation are based on Foster, et al. (*86*) (Fig. S1). Time specific solar luminosity for each simulation was based on Gough (*87*).

**Figure S1.**
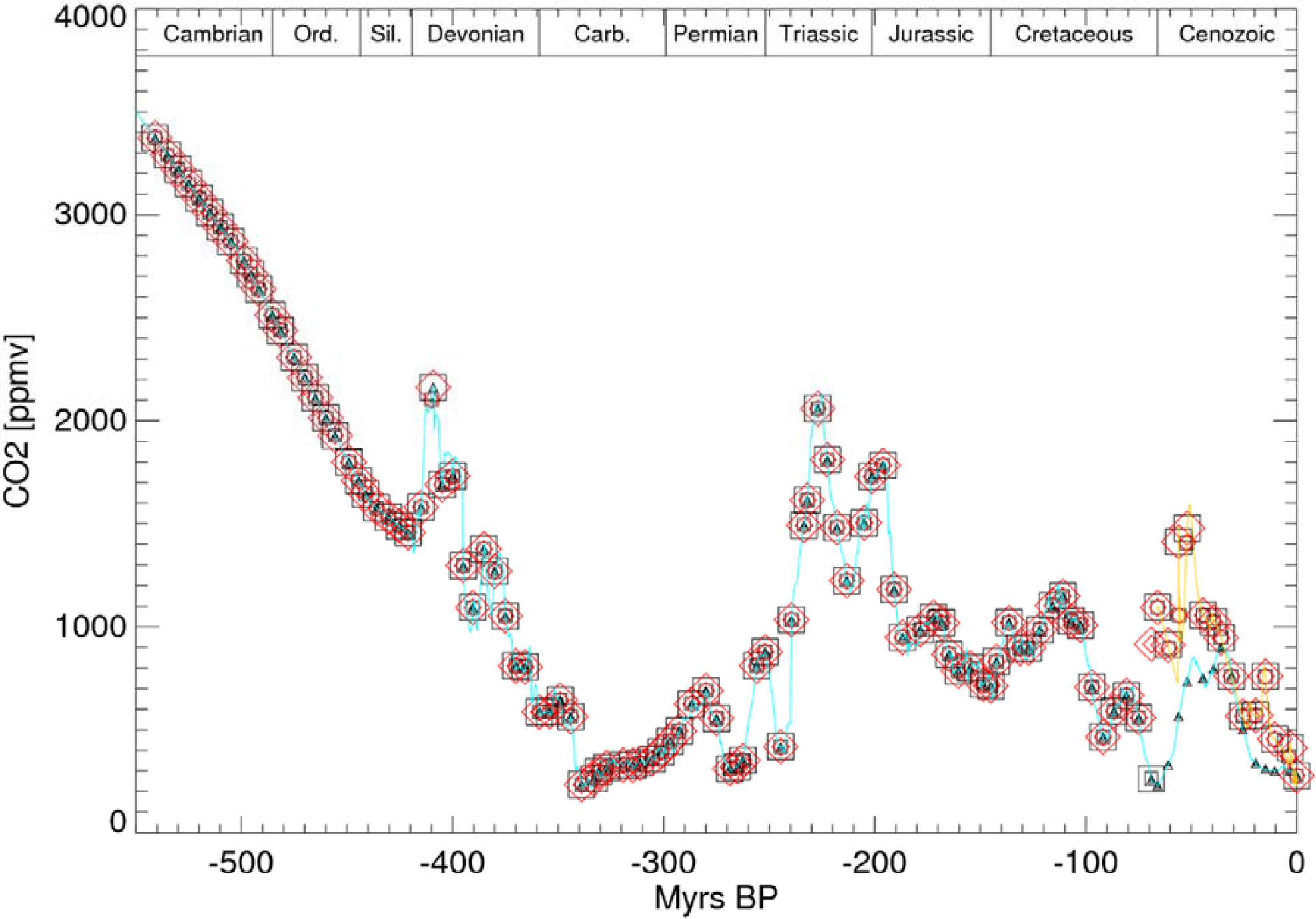
Phanerozoic (last 540 Ma) *p*CO_2_ estimates (blue line) reproduced from Foster, et al.(*86*). Red box/triangle show the target *p*CO_2_ for each snapshot simulation through the 117 individual simulations. Time is reported in the x-axis with negative numbers from the oldest dates (Cambrian, < ™500 Myrs BP) to the present (0 Myrs BP). Abbreviations: ppmv, parts per million by volume; Myrs BP, million years before present.

### Phylogenetic Principal Component Analyses

To evaluate climatic niche space occupation in a multivariate setting (combining several variables, like temperature and precipitation) we used Phylogenetic Principal Component Analyses (phyloPCA), which takes the non-independence between related species when computing covariates compared to a classic PCA (*88*). First, we checked for collinearity of variables between all the physical outputs of the GCM, including seasonal and monthly temperature and precipitation values, and retaining those variables demonstrating a Pearson’s correlation coefficient of < 0.7 (Fig. S2). The variables retained for the final phyloPCA included maximum mean annual temperature, minimum mean annual temperature, minimum mean annual precipitation and precipitation seasonality. As a phylogenetic framework, we used an updated composite supertree for all Dinosauria (Data S1) from Benson et al. (*4*, *70*) which contains 642 tips of non-dinosaurian Dinosauromorpha and the three Dinosauria subclades (Ornithischia, Sauropodomorpha and Theropoda). To maximise phylogenetic control (i.e. controlling for phylogenetic signal) in the structure of the variance in our Principal Components, we used an optimized Pagel’s λ (*89*) transformation. The object scores of the two main PCA axes (PC1 and PC2) were then extracted (Data S1) and used to model the occupation in multidimensional climatic niche space (see Macroevolutionary OU modelling section below). We used the R package phytools v.1.9-16 (*90*) for phyloPCA analysis and plotting.

**Fig S2.**
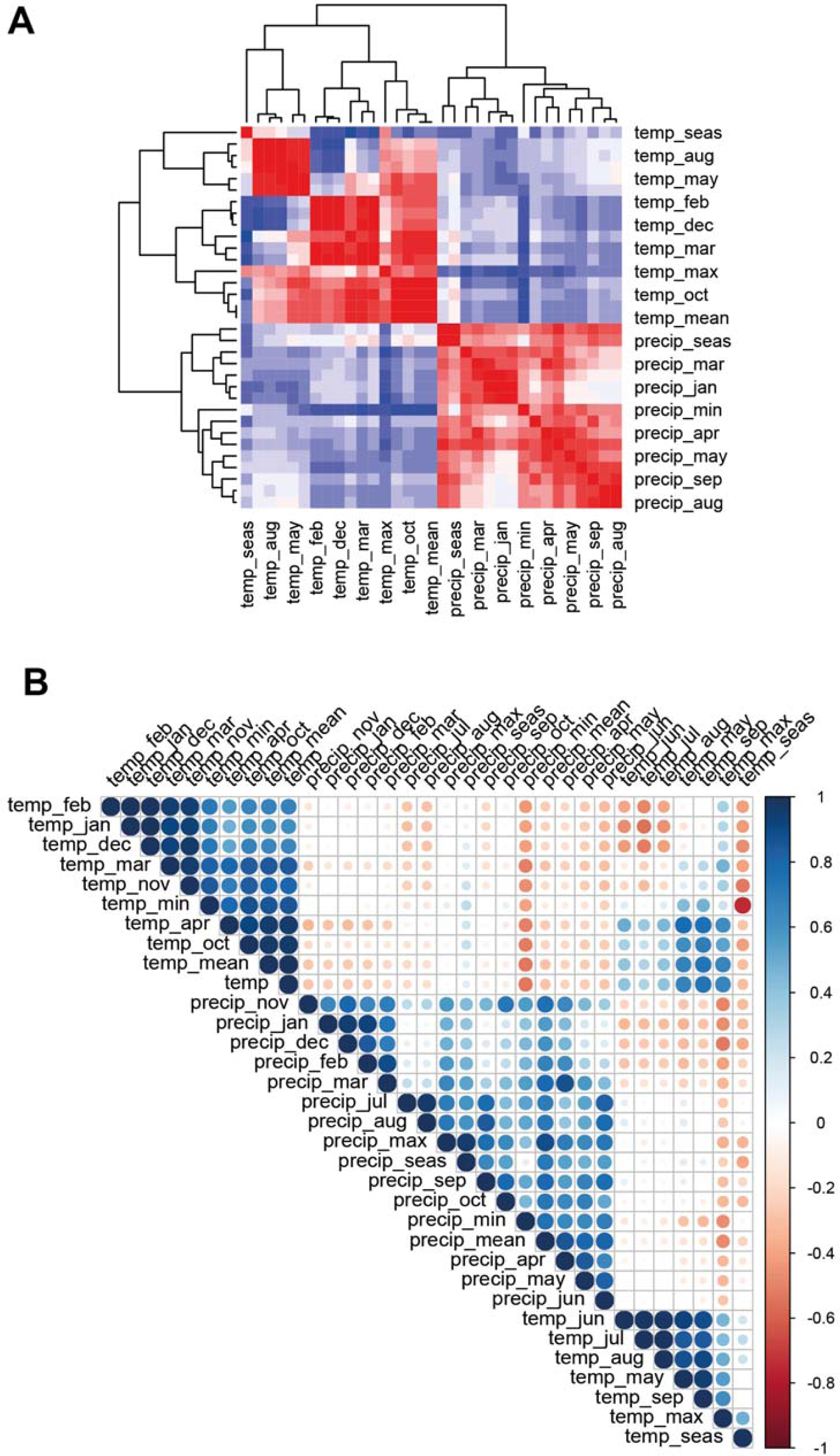
Correlograms for evaluating the collinearity between GCM output variables. Abbreviations: seas (seasonal variables), max (maximum), min (minimum), all other abbreviations are the first 3 initials of the 12 months in the modern-day Gregorian calendar.

### Phylogenetic Data for macroevolutionary modelling

In this study, we focused on the largest phylogenetic analyses currently available, closer to the current consensus on in group relationships, and including as many taxonomic tips as possible for the 3 main dinosaur subclades: Ornithischia, Sauropodomorpha and Theropoda (Data S1). A phylogeny for each subclade was sourced from Benson et al. (*4*) (originally published by Benson et al. (*70*)). Notably, the theropod phylogeny from Benson et al. (*4*) included taxa from the Carnian stage of the Upper Triassic to the Aptian stage of the Lower Cretaceous. To compensate for the lack of post-Aptian theropods, and given our interest in exploring the trends in theropod climatic niche space up to the Cretaceous/Paleogene boundary, we conducted macroevolutionary modelling analyses for Theropoda using the phylogeny published by Cau (*56*). This latter, more complete phylogeny of Theropoda, is based on a phylogenetic data matrix including 1781 morphological character statements for 132 operational taxonomic unit (*56*). Given the original criteria for assembling this dataset included, beyond broad taxonomic sampling, preferring taxa with higher amount of skeletal completeness and more robustly placed positions related to the consensus expressed by other phylogenetic analyses (*56*), the output tree from this matrix was preferred for phylogenetic comparative analyses rather than using a secondarily assembled, composite supertree.

The tree used for Sauropodomorpha (*4*) contains 98 tips (Fig. S3), the tree for Ornithischia (*4*) contains 126 tips (Fig. S4), while the tree for Theropoda (*56*) contains 132 tips (Data S1). In order to congruently fit the phylogenetic theropod tree tips with the available climatically calibrated fossil occurrences, 33 taxa were excluded from the analysis, rendering a final tree with 99 tips (Fig. S5). The tips dropped in the final tree from Cau (*56*) are: *Meleagris, Teleocrater, Euparkeria,Buriolestes, Sanjuansaurus, Chilesaurus, Bicentenaria, Gualicho, Jianchangosaurus, Halzkaraptor, Jianianhualong, Gobivenator, Anchiornis* (holotype), *Serikornis, Aurornis, Cruralispennia, Chongmingia, Zhouornis, Sulcavis, Bohaiornis, Parahongshanornis, Archaeornithura, Piscivoravis, Iteravis, Yi, Berberosaurus, Coelurus, Eodromaeus, Epidendrosaurus, Guaibasaurus, Shenzhouraptor, Sinocalliopteryx* and *Sinusonasus*.

To deal with phylogenetic uncertainty related to different topologies (due to slightly inconsistent inter-nodal relationships in each of the phylogenies of the selected subclades), we randomly chose one of the 20 trees in each tree object for the Benson et al. dataset (*4*). The Cau (*56*) theropod tree used is the Maximum Clade Credibility Tree topology (MCCT) obtained as output of the Bayesian analyses (see details in Cau (*56*)). Time calibration for the Benson et al. dataset (*4*) was performed by assigning ages in each tip as the uniform distributions between the minimum (LAD) and maximum (FAD) possible ages for each taxon, with both FAD and LAD being scrutinised, vetted and updated for this study (see Data S1), and with node age calibration following the Lloyd et al. (*91*) modified probabilistic method by Hedman (*92*). These trees (*4*) were calibrated with their roots at 239.7856 Ma while the more inclusive, Cau (*56*) tree was calibrated with the root at 252 Ma.

**Fig. S3.**
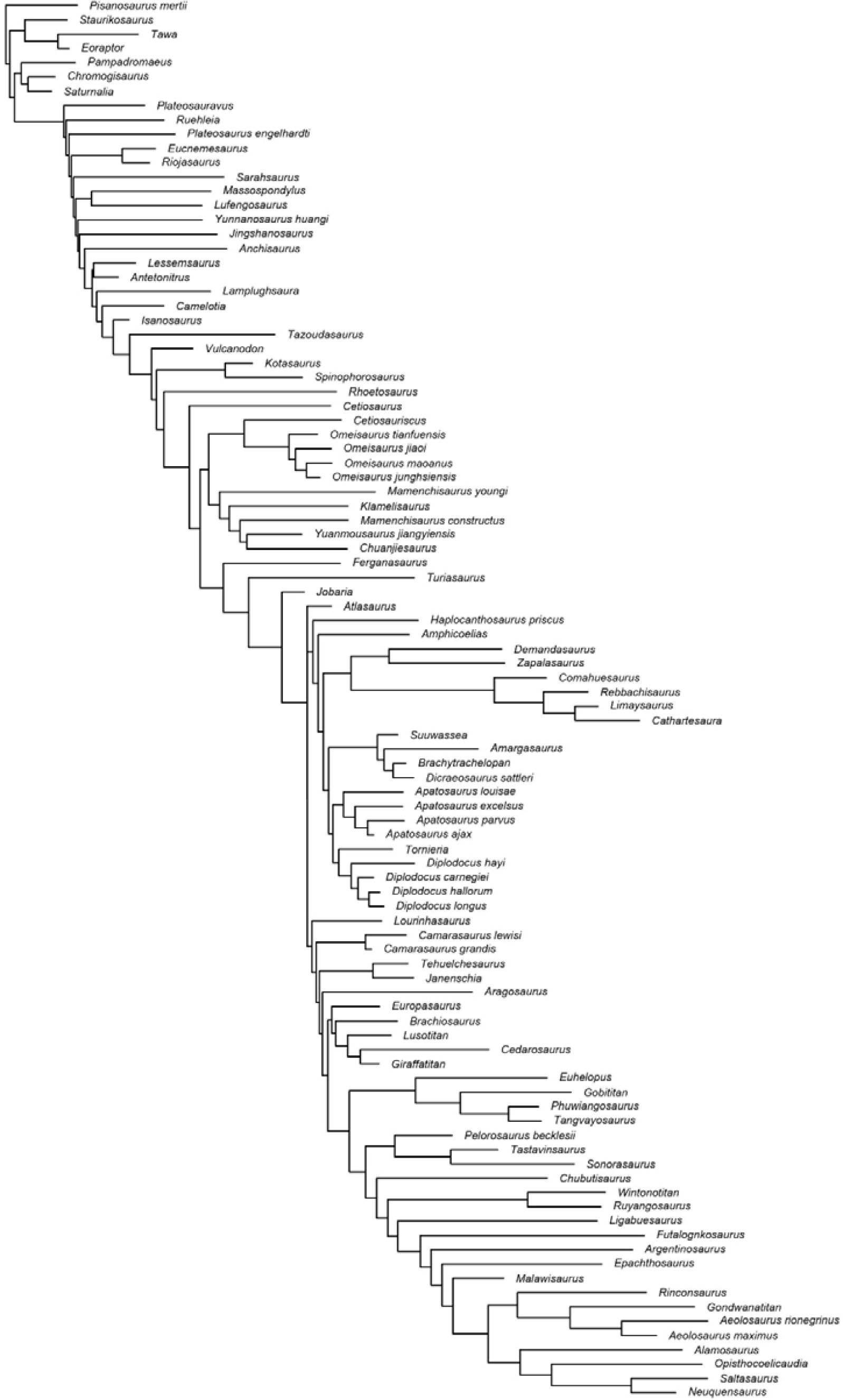
Phylogenetic tree of Sauropodomorpha used in this study.

**Fig. S4.**
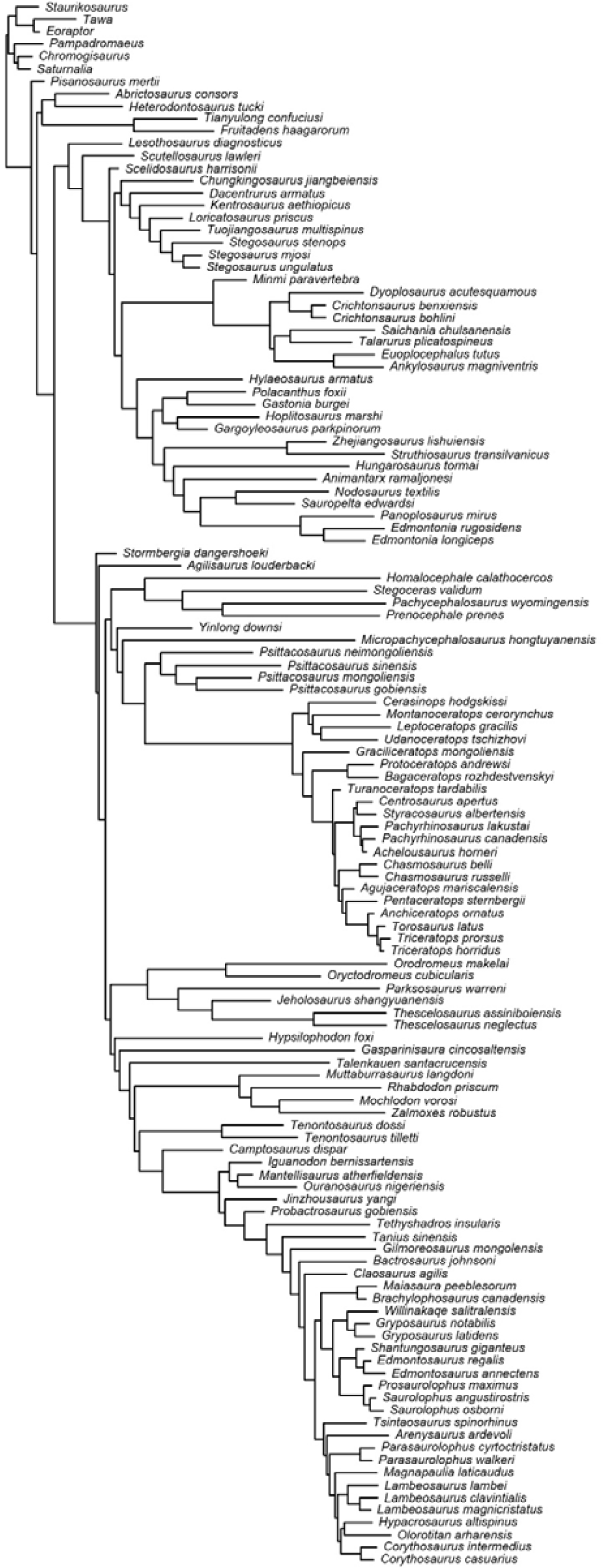
Phylogenetic tree of Ornithischia used in this study.

**Fig. S5.**
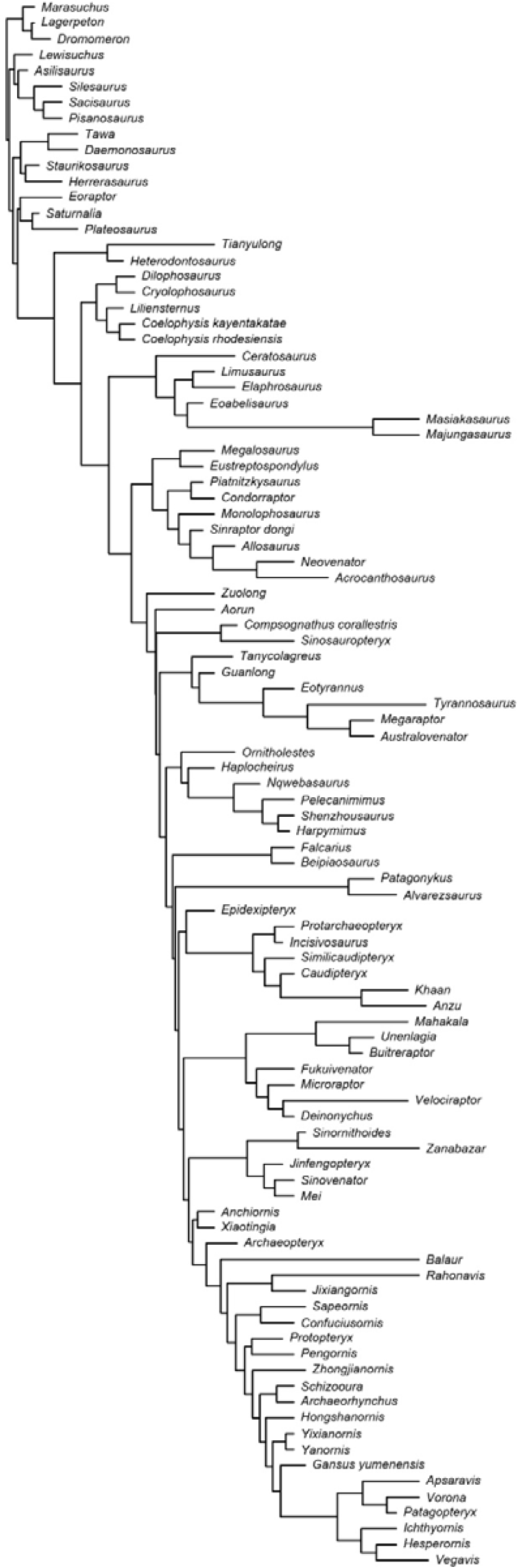
Phylogenetic tree of Theropoda used in this study.

### Ancestral State Reconstruction

We performed ancestral state reconstruction to map the evolution of physical environmental parameters, such as temperature, precipitation, and principal components derived from multidimensional climatic space (from phyloPCA, see Phylogenetic Principal Component Analyses section), treating them as continuous characters. We used the R package phytools v.1.9-16 (*90*) to create a stochastic map with 10,000 generations, applying the ‘SYM’ (Symmetrical) model of evolution on the randomly selected, time-scaled consensus tree, following the approach by Gates et al. (*93*) as used in Chiarenza et al. (*65*) to reconstruct body-size evolution in ornithischian dinosaurs. The resulting ancestral state reconstruction was visualized as a density map on the phylogenetic trees for each dinosaur subclades (Fig. S6–S11). Furthermore, we utilized a GEIGER- fitted comparative model for continuous data (*94*) to reconstruct the ancestral state (z_0_ or root value) at the base of various dinosaur subclades Fig. 2. For Ornithischia, ancestral states were reconstructed at the tree root (*Staurikosaurus*+*Fruitadens haagarorum*), for Thyreophora (*95*–*97*) (*Scutellosaurus lawleri*+*Edmontonia*), and for Neornithischia (*95*, *96*, *98*) (*Stormbergia dangershoeki*+*Corythosaurus*). Ancestral States Reconstructions (ASRs) were produced for Sauropodomorpha at the tree root (*Pisanosaurus*+*Chuanjiesaurus*), for non-somphospondylian Neosauropoda (*52*, *99*) (*Ferganasaurus*+*Giraffatitan*), and for Somphospondyli (*100*, *101*) (*Euhelopus*+*Saltasaurus*). ASRs were computed for Theropoda (*56*), testing two different sets of ASR partitions: one (RTC) at the root of the tree including non-theropod dinosauromorphs (*Marasuchus*+*Heterodontosaurus*), non-coelurosaurian Theropoda (*95*, *102*) (*Dilophosaurus*+*Acrocanthosaurus*), and Coelurosauria (*102*) (*Zuolong*+*Vegavis*); the other regime partition (RCA) included the root of the tree with non-coelurosaurian dinosauromorphs (*Marasuchus*+*Acrocanthosaurus*), for non-avialan Coelurosauria (*102*–*104*) (*Zuolong*+*Mei*), and for Avialae (*102*) (*Anchiornis*+*Vegavis*).

**Fig. S6.**
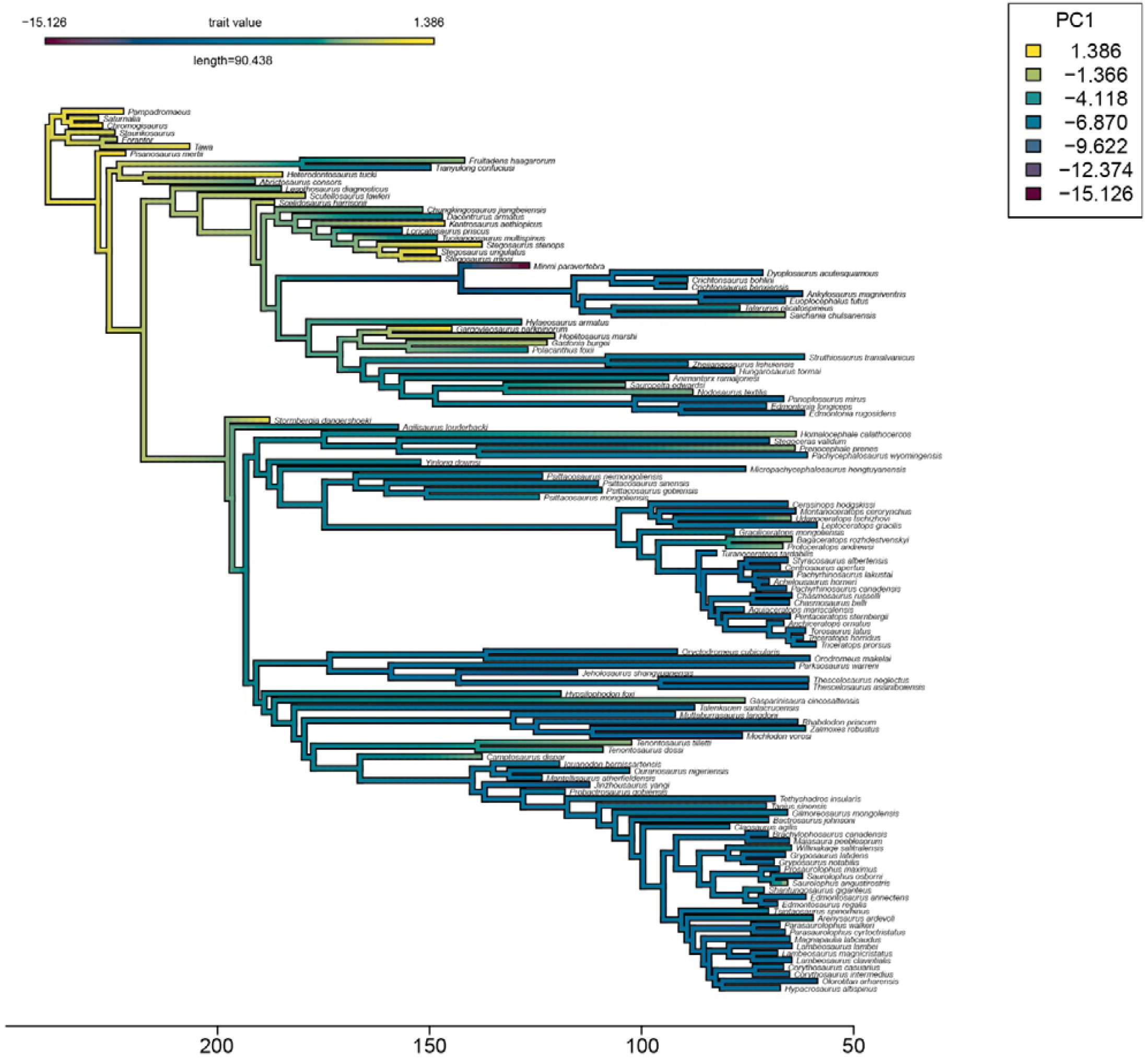
Ancestral state reconstruction for Ornithischia phylogeny for the PC1 trait.

**Fig. S7.**
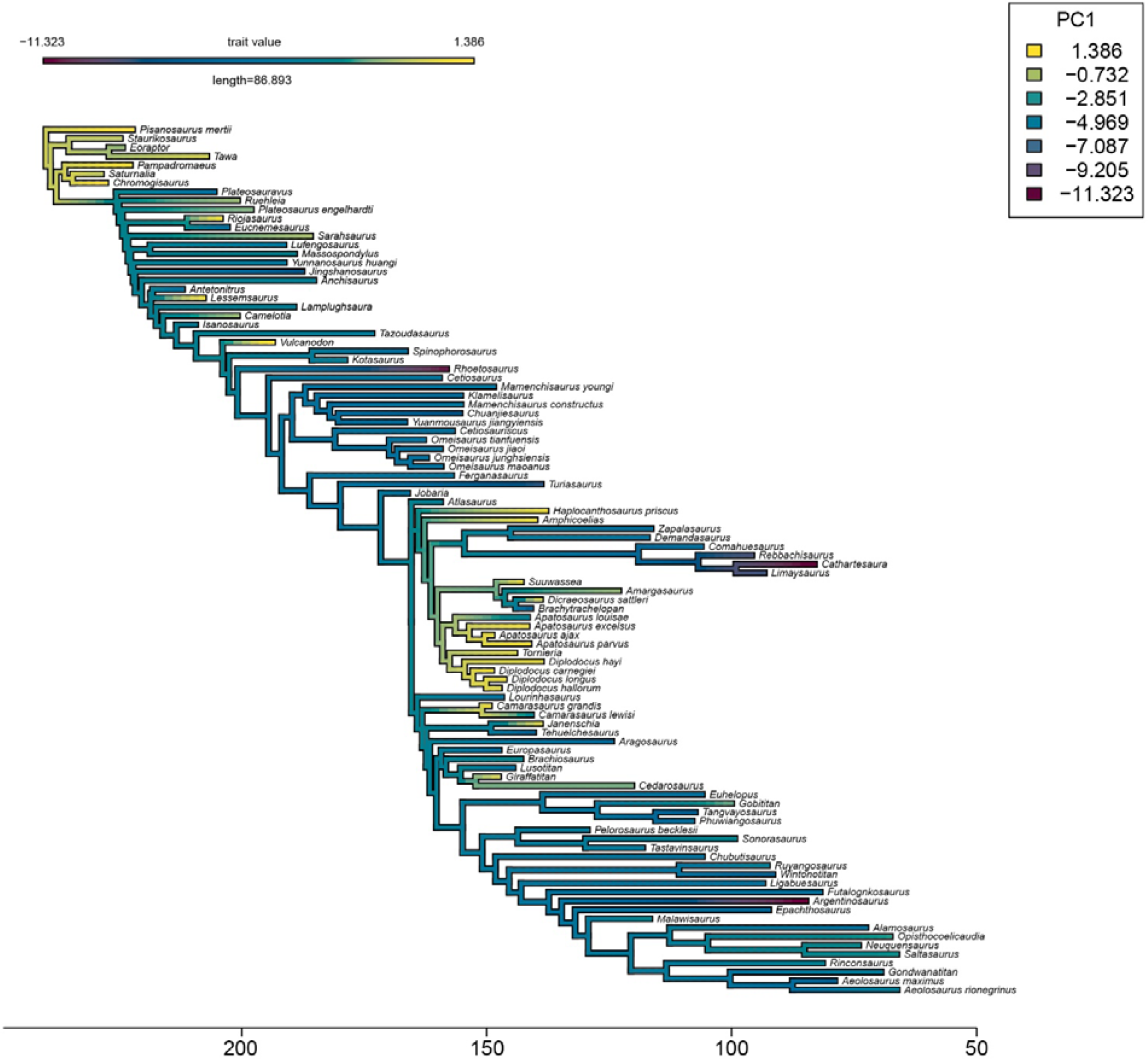
Ancestral state reconstruction for Sauropodomorpha phylogeny for the PC1 trait.

**Fig. S8.**
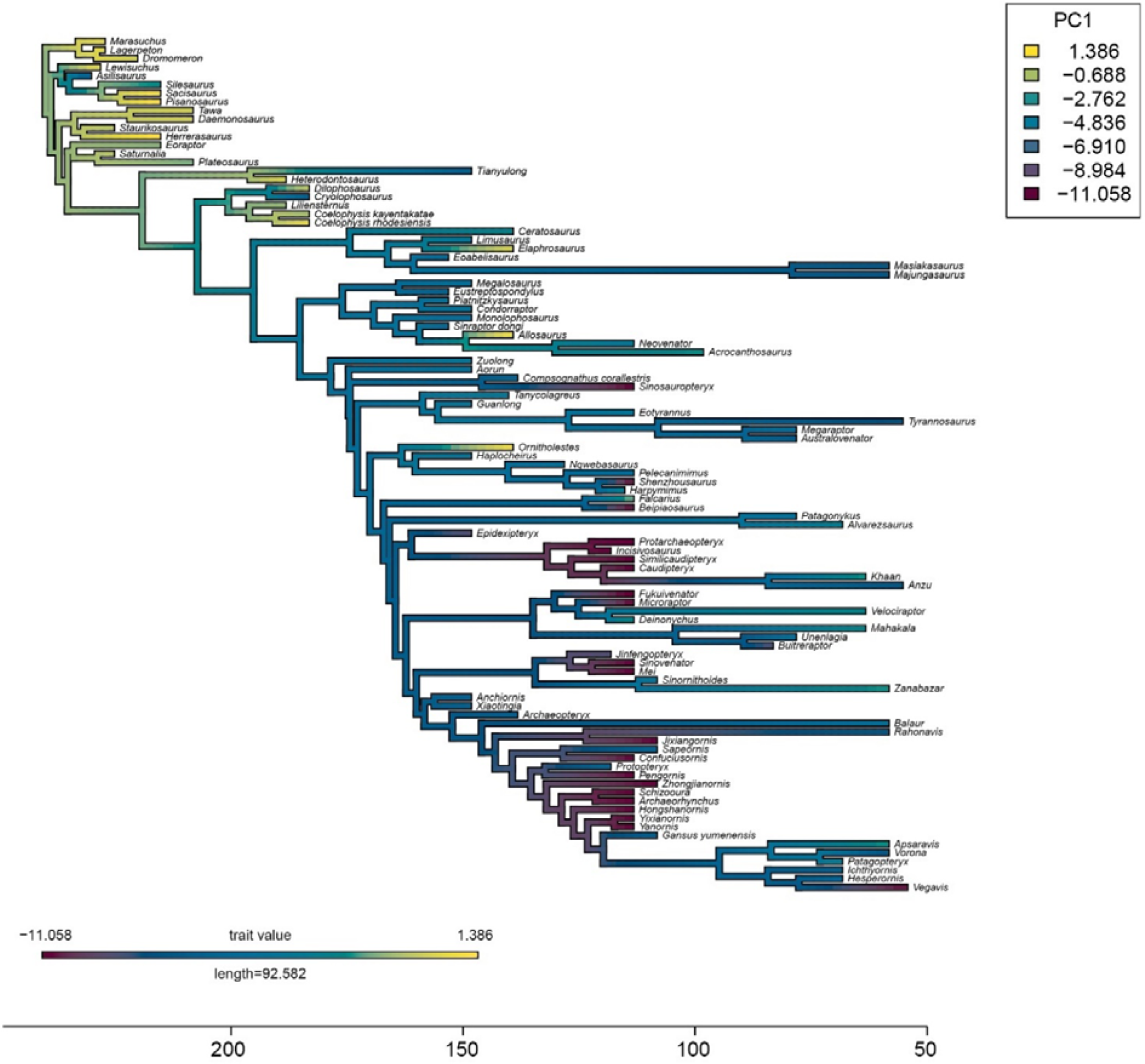
Ancestral state reconstruction for Theropoda phylogeny for the PC1 trait.

**Fig. S9.**
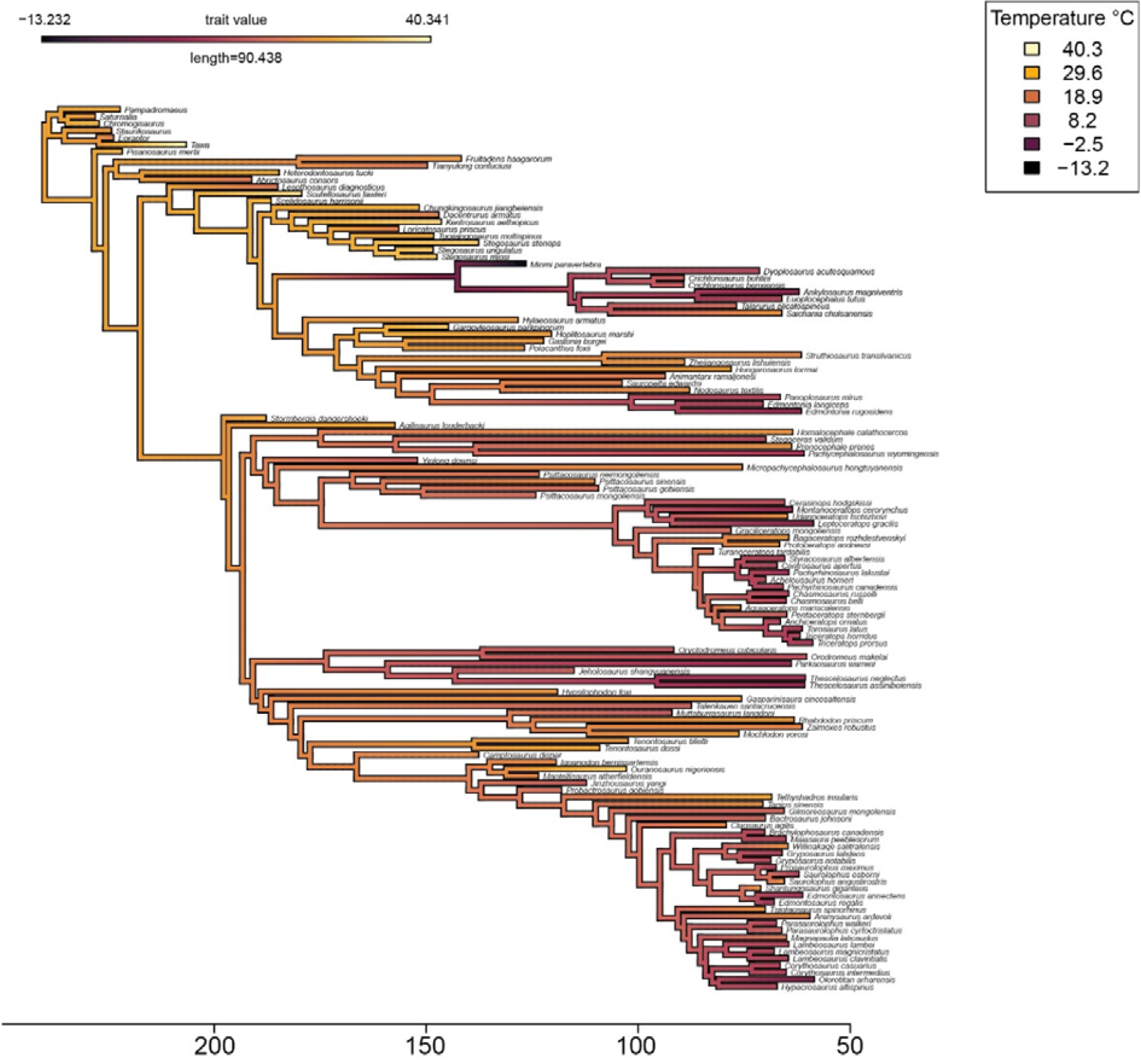
Ancestral state reconstruction for Ornithischia phylogeny for the temperature trait.

**Fig. S10.**
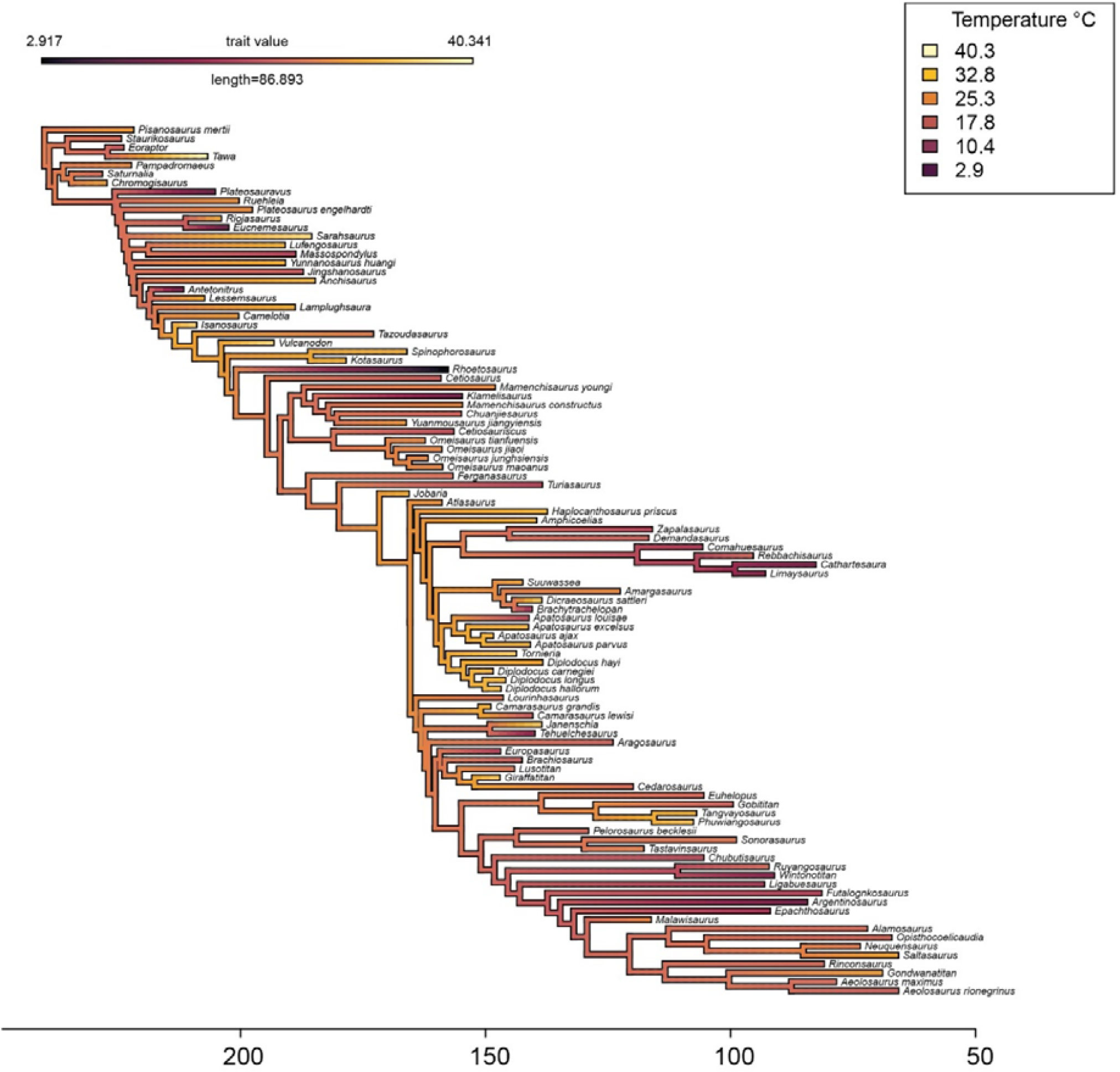
Ancestral state reconstruction for Sauropodomorpha phylogeny for the temperature trait.

**Fig. S11.**
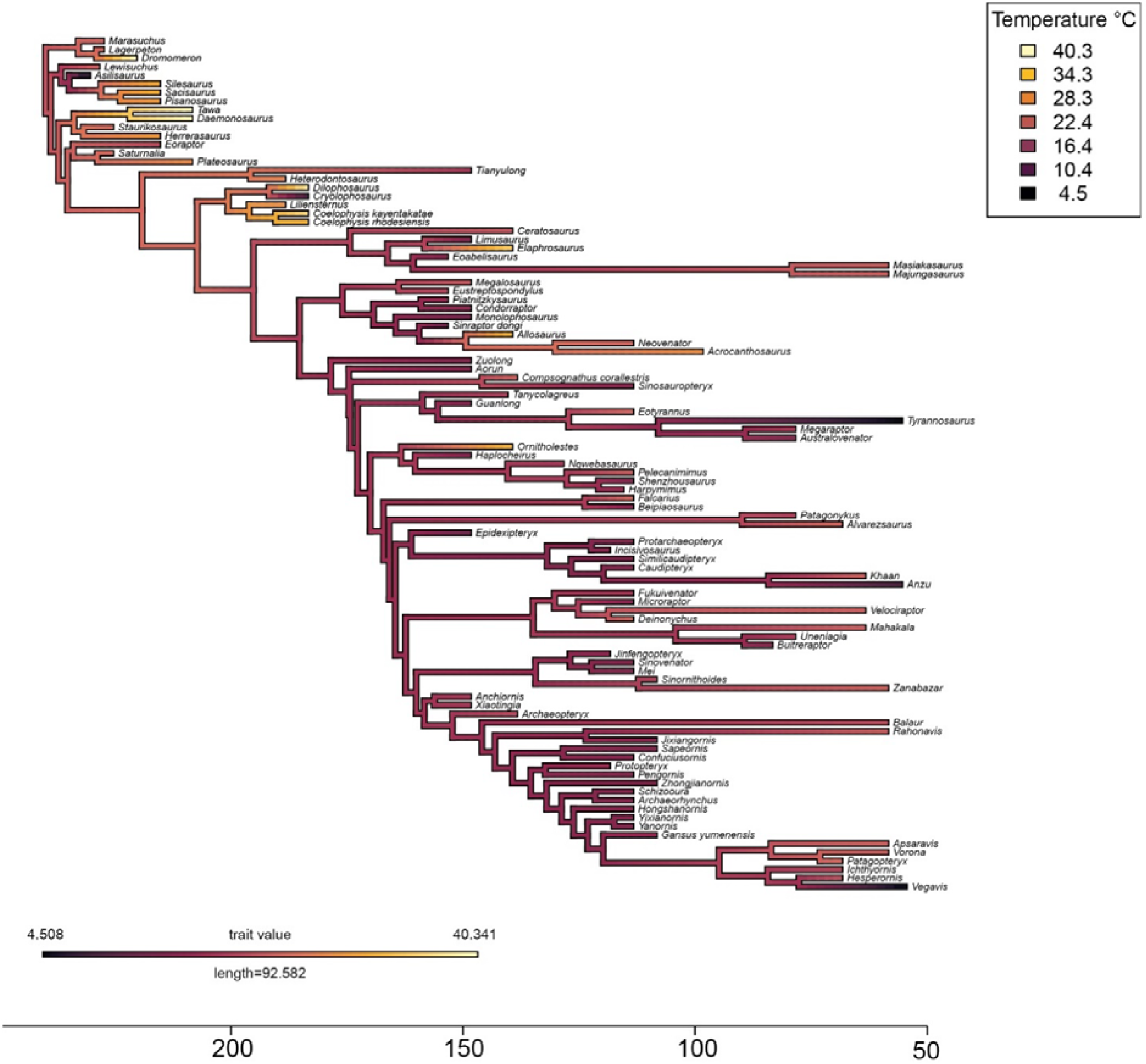
Ancestral state reconstruction for Theropoda phylogeny for the temperature trait.

### Macroevolutionary OU modelling

We explored the evolution of the climatic landscape (niche space) of each dinosaur subgroup by means of Ornstein–Uhlenbeck (OU) dynamics (*34*, *35*, *105*). Using this approach, we aimed to test the congruence of dinosaurian climatic landscape exploration against several constrained and well-documented evolutionary scenarios (like Brownian motion (*106*), Early burst (*107*), and multiDpeak OU (*34*) models of evolution).

To accomplish this, we initially calibrated the phylogenetic tree with co-occurring climatic variables for the specimen-based tips (OTUs, Operation Taxonomic Units). Subsequently, we performed macroevolutionary modelling using OU models to investigate whether the evolution of each specific climatic trait value, followed a stochastic diffusion pattern over time (like in a Brownian motion (*106*) scenario) or a directional, trend-like occupation (OU models) of climatic niche space. Given the predominance of temperature in explaining the variance of the data (see phyloPCA Methods section and Fig. 1), and the higher reliability (higher ‘realism’) of temperature as a GCM output compared to precipitation variables (*80*), we further discussed and plotted only temperature related macroevolutionary models in the main results. We assessed whether each phylogenetic model followed any direction trend toward a macroevolutionary trait optimum (θ) at a given rate of variance accumulation (σ), with a defined strength of attraction (α) at a particular phylogenetic node (OU model), at multiple nodes (OUM multiple regimes (*34*, *35*, *105*)) or under different time-partitions (Data S3). Temporal partitions (Data S2) for each tree were performed at the Jenkyns event (*58*, *108*) (early Toarcian, 183.0 ma), at the Jurassic/Cretaceous boundary (*109*) (145.0 Ma) and at the Cenomanian/Turonian boundary (*57*, *110*) (93.9 ma).

Model performance was compared (Tables S1, Data S3) by using the Akaike Information Criterion with finite correction (*111*), excluding models that returned out of bounds (α, σ, θ) estimates. For the analysis of continuous trait evolution under selective regimes, we employed modified OU models (*34*), using R version 4.3.0 and the OUwie package v.2.10 (*112*) with codes from Revell and Harmon (*113*) (see also http://www.phytools.org/Rbook/).

**Table S1.** Model selection based on AICc. This table shows a selection of the first 10 best performing models for each experiment. For complete table with evaluation of all the models see Data S3. Models italicized and highlighted in bold are the best performing for both temporal and clade partitions in each clade. Abbreviations for models of trait evolution: BM, Brownian motion; clade, models with different parameters for each clade; EB, early burst (high σ at the base of the tree); temporal, a model with a significant shift at a given time slice, here selected at 183 million years ago (Jen, ‘Jenkyns’ event), 145 million years ago (J/K, Jurassic/Cretaceous boundary) and at 93.9 million years ago (C/T, Cenomanian/Turonian boundary); Stasis, a model with strong α towards a given θ; OU, Ornstein–Uhlenbeck; OUM, multiDpeak OU model (multiple regimes within individual θ values) with fixed values of α and σ; OUMV, multiDpeak OU model with fixed values of α and σ as a free parameter; OUMA, multiDpeak OU model with fixed values of σ and α as a free parameter; OUMVA, multiDpeak OU model with both σ and α as free parameters; Trend, a model following a given linear trend (given α constant). Clade-partitioned models in Theropoda are abbreviated as: RTC, three portioned model with Root, Theropoda and Coelurosauria; RCA, three partitioned model with Root, Coelurosauria and Avialae.

| Ornithischia | AICc | Sauropodomorpha | AICc | Theropoda | AICc |
| --- | --- | --- | --- | --- | --- |
| ***OUMVA temporal CT*** | -784.9 | ***OUMA temporal Jen*** | -622.5 | ***OUMVA temporal Jen*** | -639.1 |
| OUMVA temporal Jen | -784.7 | OUMA temporal JK | -621.0 | ***OUMVA clade RCA*** | -631.3 |
| OUMA temporal JK | -783.0 | OUMVA temporal Jen | -620.7 | OUMVA clade RTC | -631.0 |
| OUMVA temporal JK | -780.7 | OUMVA temporal JK | -617.4 | BMS clade RCA | -614.8 |
| ***OUMA clade*** | -776.6 | OUMA temporal CT | -615.8 | BMS clade RTC | -613.8 |
| OUMVA clade | -776.0 | OUMVA temporal CT | -615.4 | OUMV clade RCA | -608.7 |
| OUM temporal CT | -760.5 | ***OUMVA clade*** | -613.4 | BM temporal Jen | -608.4 |
| OUM temporal Jen | -760.1 | BMS clade | -597.0 | OUMV clade RTC | -607.9 |
| OUMV temporal CT | -759.9 | OUMV clade | -594.1 | OUMV temporal Jen | -605.4 |
| OUMV temporal Jen | -759.9 | eb clade | -589.6 | eb Theropoda RTC | -602.1 |

### Climatic zones

Using our HadCM3L paleoclimatic simulations (see related ‘Paleoclimate model’ section above), we classified landscapes into broad climatic zones following an adapted version of Köppen’s climate classification. Retaining temperature and precipitation monthly data per time step (formatted as two arrays of twelve matrices, respectively), we implemented (*39*) the Köppen-Geiger(*38*) climate classification from Beck et al. (*114*). To do so, we used the provided function written in MATLAB (https://doi.org/10.6084/m9.figshare.6396959) to classify these climate data into five main climate classes: tropical, arid, temperate, cold, and polar. We introduced two changes to the original Beck et al. (*114*) function: i) we added a “-99” value for the missing values, in order to facilitate computational operations, and ii) we modified lines 84, 85 and 86 of KoppenGeiger.m in order to fulfill the following conditions: “Pthreshold =2×MAT if >70% of precipitation falls in winter, Pthreshold =2×MAT+28 if >70% of precipitation falls in summer, otherwise defined as Pthreshold =2×MAT+14” (H. Beck, personal communication, December 18, 2021). Once we have obtained the classified matrices per time step, we rasterized and rotated the resulting maps using the ‘rast’ and ‘rotate’ functions from the terra package version 1.7-39 (*115*).

We downloaded land configuration maps for all time periods using the ‘getmap’ function from the mapast package version 0.1 (*116*), using the PALEOMAP (*117*) reconstruction model as it is the same used as boundary conditions for our climate data. We cropped the previous reclassified maps with these paleogeographic maps using the ‘mask’ function of the R package terra (version 1.7-39) (*115*), to retrieve land climatic zones’ maps. Lastly, we used dinosaur paleolatitude and paleolongitude from our dataset (Data S1) to extract climatic zone information from each of the occurrences (Fig. S12–S14). To do so, we used the ‘extract’ function of the terra version 1.7-39 package (*115*). With this information, we calculated the number of dinosaur occurrences by subclade included in our trees (Ornithischia, Sauropodomorpha and Theropoda) and by climatic zone through time to obtain the quantification shown in Figure 3.

**Fig. S12.**
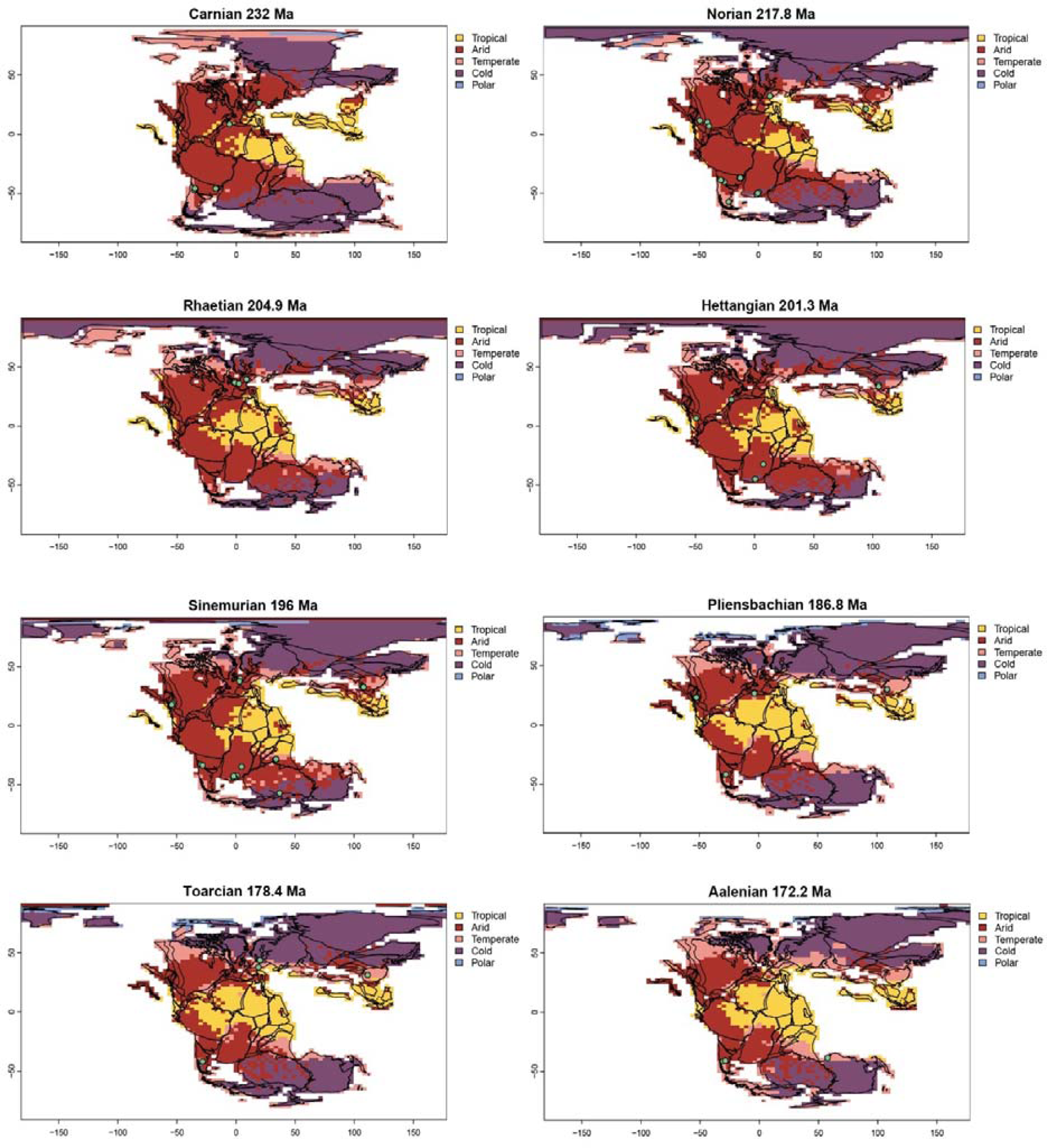
Paleogeographic maps for the Carnian–Aalenian interval (232–172.2 million years ago) showing climatic zones modelled in this study from paleoclimate data. Abbreviation: Ma, mega annum (million years ago). Green dots representing dinosaur bearing localities for taxa in our phylogenetic trees.

**Fig. S13.**
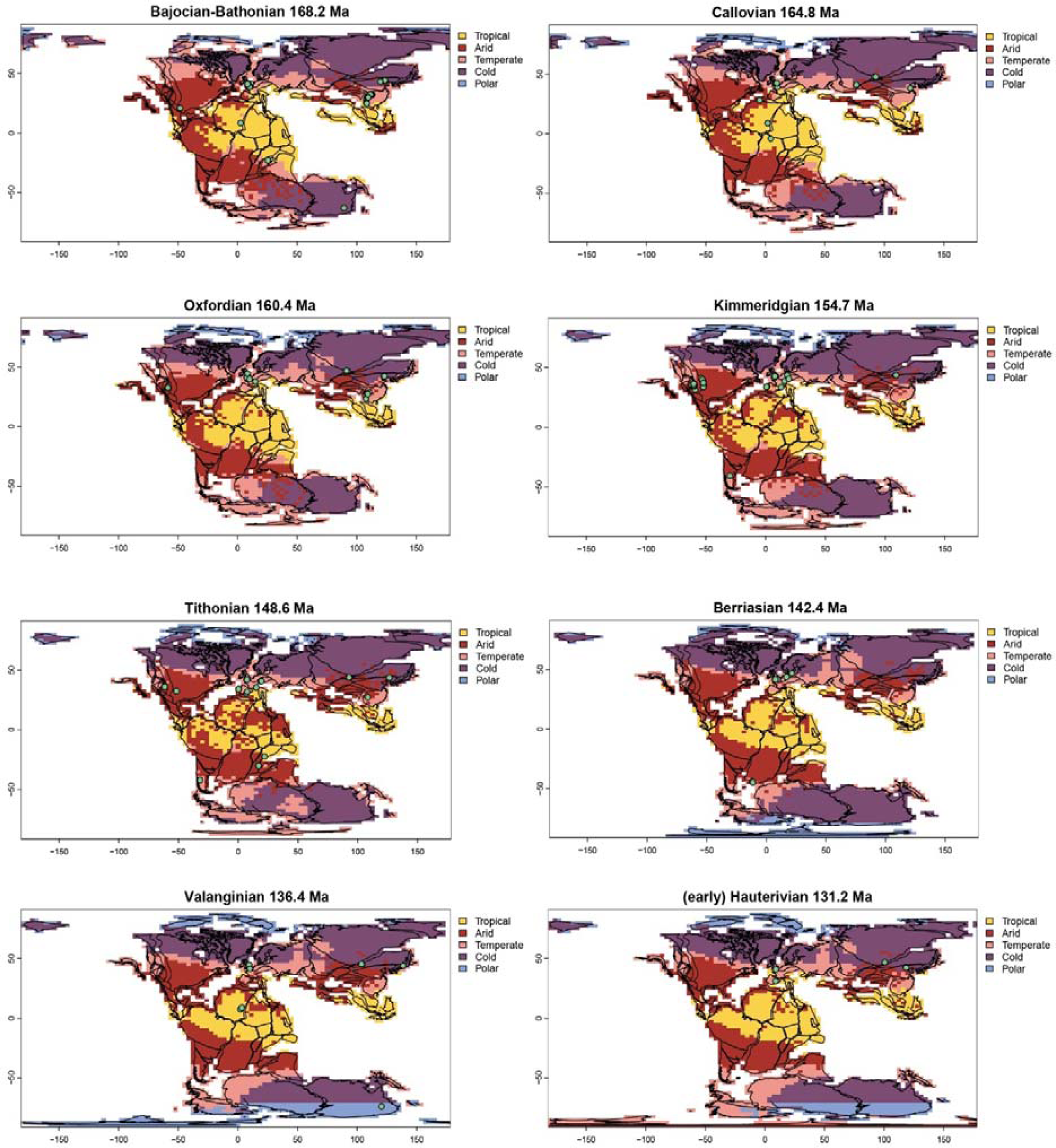
Paleogeographic maps for the Bajocian–(early) Hauterivian interval (168.2–131.2 million years ago) showing climatic zones modelled in this study from paleoclimate data. Abbreviation: Ma, mega annum (million years ago). Green dots representing dinosaur bearing localities for taxa in our phylogenetic trees.

**Fig. S14.**
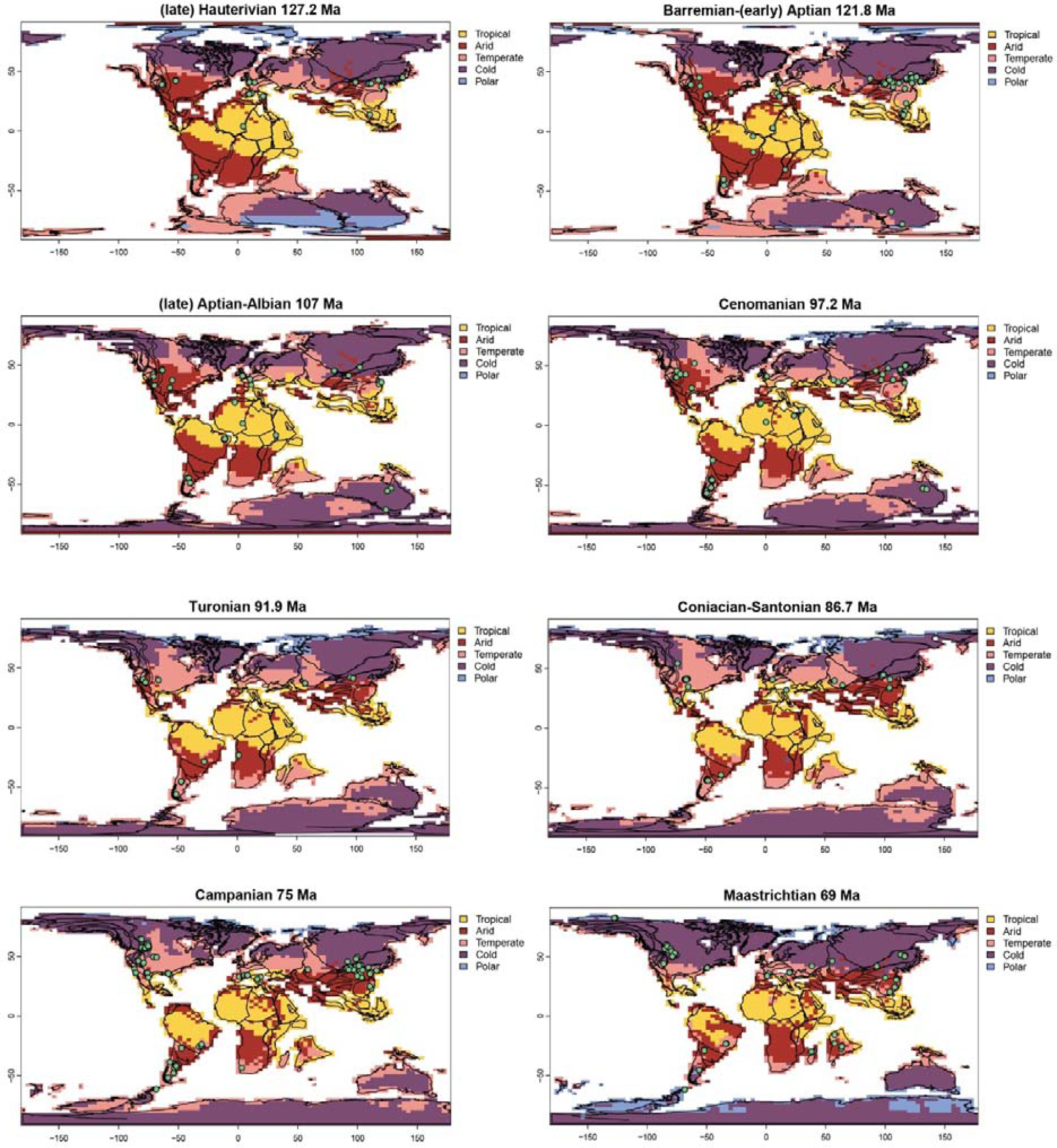
Paleogeographic maps for the (late) Hauterivian–Maastrichtian interval (127.2–69 million years ago) showing climatic zones modelled in this study from paleoclimate data. Abbreviation: Ma, mega annum (million years ago). Green dots representing dinosaur bearing localities for taxa in our phylogenetic trees.

